# AtlasOT - The Fused Unbalanced Gromov-Wasserstein for Multimodal Integration of Disease Atlases

**DOI:** 10.64898/2026.09.22.753496

**Authors:** Kai Peng, Mayra Ruiz, Barthélémy Caron, Christoph Kuppe, James S. Nagai, Ivan G. Costa

## Abstract

Single-cell and spatial multiomics technologies are transforming disease atlas construction, but computational integration of unpaired modalities remains challenging, as existing methods overlook two biological constraints of data with matching samples. First, cells should only be mapped across modalities within the same donor or biospecimen, and second, cell recovery frequently differs substantially between modalities. To address these gaps, we present AtlasOT, an optimal-transport framework based on the fused unbalanced Gromov-Wasserstein (FUGW) formulation for multi-modal integration. AtlasOT jointly models a shared cross-modality feature space and modality-specific geometric structures while restricting transport to within-sample cell pairs. Moreover, AtlasOT relaxes strict mass conservation of the balanced optimal transport optimization to accommodate unbalanced cell numbers. We benchmark AtlasOT against state-of-the-art methods on scRNA-scATAC and scRNA-spatial transcriptomics mapping scenarios. Our results indicate that AtlasOT outperforms baselines and state-of-the-art methods in all considered scenarios. Moreover, we demonstrate that AtlasOT’s transport plan can be used to improve several relevant tasks such as the detection of rare cell populations, spot-level cell-type deconvolution, spatial gene imputation, and reconstruction of transcription-factor-driven spatial regulatory dynamics. These results establish AtlasOT as a unified, biologically constrained framework for multimodal integration in disease atlas studies, with broad applicability to label transfer, imputation, and regulatory network analysis.

## Introduction

Single-cell and spatial transcriptomics sequencing are disruptive technologies allowing the quantification of transcriptional status (scRNA-seq) or open chromatin status (scATAC-seq) of dissociated single cells^1,2^ or intact tissues (spatial transcriptomics; ST)^3^. The decrease in sequencing prices is making the measurement of multimodal single-cell disease atlases possible, capturing hundreds of thousands of cells across distinct individuals and modalities, as in the spatial heart atlas of myocardial infarction^4^ or the kidney precision medicine project^5^. While novel protocols allow joint profiling of RNA and open chromatin in single cells or spatial locations^6,7^, technical limitations, including low cellular recovery, limit their adoption. Existing multimodal disease atlases mostly rely on measurements of consecutive cross-sections of the same biological specimen.

Several computational approaches for integrating scRNA-seq, scATAC-seq, and spatial transcriptomics have been proposed, aiming to overcome modality heterogeneity, data sparsity, and the lack of paired measurements^8,9^. Among others, challenges include the fact that each modality might have distinct feature spaces (genes for scRNA-seq vs peaks for scATAC-seq) and different resolutions (single-cell vs spatial spots containing several cells). There is a large computational and benchmarking literature on the integration of unpaired single-cell data (diagonal integration)^10–13^ and single-cell to spatial mapping^14^. However, to the best of our knowledge, no study considers the fact that cells in a disease atlas should not be mapped across samples or biospecimens, as this could lead to a mapping where a cell from a disease sample can be mapped to a cell from a healthy sample. Another overlooked aspect is the fact that cell recovery differs between protocols^15,16^. For example, we could recover three times more scRNA-seq cells than scATAC-seq cells from consecutive cross-sections, despite the use of the same isolation protocol^4^. These important points have been overlooked by all previous benchmarking studies^10–12^, as they rely on balanced paired multimodal data, i.e., they have the same number of scRNA-seq and scATAC-seq cells. In short, state-of-the-art methods are evaluated under simple and unrealistic benchmarking scenarios, as real-world non-paired datasets are imbalanced.

Recently, optimal transport (OT)-based methods have drawn attention for their ability to estimate mappings between scRNA-seq and scATAC-seq^17,18^ or between scRNA-seq and spatial transcriptomics^19,20^. These methods explored variants of OT algorithms, such as Gromov-Wasserstein^17,19^ or Fused Gromov-Wasserstein algorithms^18^, which allow every modality to be represented in a distinct feature/metric space. Also, unbalanced optimal transport has been proposed to account for the fact that the distribution of cells can differ between the two modalities^21^. However, none of the previous approaches considers the fact that mapping should be constrained to cells from the same biospecimen or donor.

## Results

### Overview of AtlasOT

To integrate multimodal disease atlases, we need to map cells across modalities — for example, linking an RNA profile to its corresponding chromatin profile, or placing a dissociated cell within its spatial context. Several constraints govern this mapping at the sample level of disease atlas datasets. First, cells should only be matched to others from the same donor or tissue sample, since mapping across biospecimens (e.g., disease to healthy) is not biologically meaningful. Second, cell recovery differs substantially between modalities, so the matching must tolerate unbalanced cell numbers rather than assume one-to-one correspondence. A further complication is that scRNA-seq, scATAC-seq, and spatial transcriptomics are measured in distinct feature spaces: genes for scRNA-seq, accessible chromatin peaks for scATAC-seq, and a limited, spatially resolved gene set for spatial data. These feature spaces are not directly comparable, so integration requires first establishing a shared representation before cells can be matched.

To address these, we propose AtlasOT (Atlas alignment with Optimal Transport), a unified optimal-transport-based framework for unpaired multimodal integration of disease atlas datasets. AtlasOT is composed of three main components. First, for a pair of modalities (scRNA-scATAC or scRNA-ST), it creates modality-specific spaces (source and target)^1^ and a shared space combining cells of both modalities using common genes for scRNA-ST mapping, or gene activity scores derived from peak accessibility for scRNA-scATAC (Fig. 1b). The modality-specific spaces are based on *k*-nn graphs, which focus on capturing geometric relationships within each modality^22^. Second, a cell mapping is performed via a Fused Unbalanced Gromov-Wasserstein (FUGW) formulation^23^, combining Optimal Transport (OT) for the shared-space alignment with Gromov-Wasserstein (GW)^24^ for the modality-specific structure (Fig. 1c). While the shared and modality-specific spaces are computed jointly across all samples, the mapping is performed for each sample independently, enforcing sample-specific matching. Moreover, the unbalanced formulation^25^ relaxes mass-conservation constraints to accommodate differing cell numbers between modalities. Finally, the output is a transport matrix **T** describing the mapping probability between cells across modalities, which forms the basis for downstream analyses including label transfer, shared-space inference, cell deconvolution, feature imputation and transcription factor chromatin flows (Fig. 1d).

**Figure 1.**
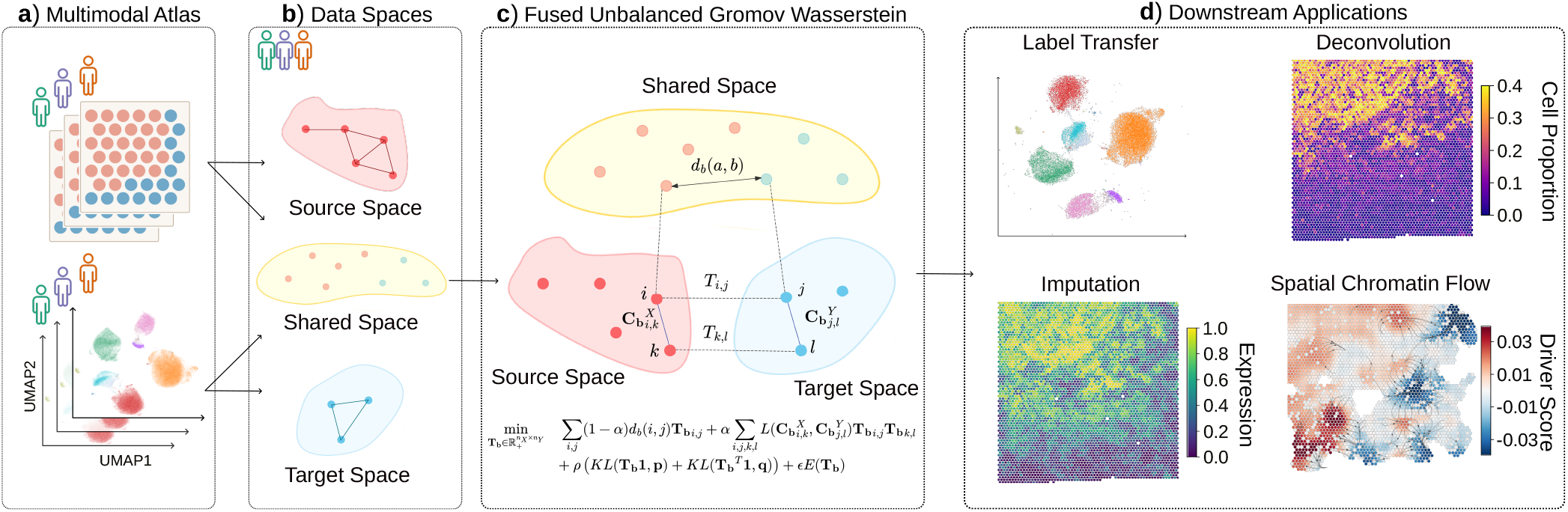
a) AtlasOT takes heterogeneous modalities—including single-cell RNA sequencing (scRNA-seq), single-cell ATAC sequencing (scATAC-seq), and spatial transcriptomics—collected from an individual biological sample as source and target domains. The framework is specifically tailored to align modalities within the same sample. **b)** AtlasOT constructs modality-specific *k*NN graphs within each individual sample to capture distinct continuous topologies and builds a shared space for cross-modal alignment. **c)** Mathematical formulation of AtlasOT solver. The FUGW objective function concurrently minimizes the cross-modal feature distances and intra-modality topological discrepancies for sample-specific integration under unbalanced marginal constraints. **d)** The optimized transport matrix is applied to facilitate downstream analyses, including cross-modal label transfer, spatial gene imputation, and spatial gene regulatory network (GRN) reconstruction.

### Benchmarking of scRNA-scATAC mapping in disease atlas

We first systematically evaluated AtlasOT and competing approaches for scRNA-seq and scATAC-seq integration (diagonal integration) across three distinct disease atlases (Muto-2021^26^, Wilson-2022^27^, Kuppe-2022^4^) totalling 38 samples and approximately 280,000 cells (Table 1). These data arise from unpaired single cell experiments, which respectively display 1.2x, 2x and 3.5x more cells from one particular modality, supporting their modality-specific imbalance. As a case study, we focus here on a label transfer problem, i.e., we use the mapping to transfer the label of the RNA-seq modality towards the ATAC-seq modality.

**Table 1.** Summary of the datasets employed in this comprehensive study across multiple biological systems and tissues.

| Dataset | Tissue | Number of cells |  | Samples | Multiome | Cell types |  |
| --- | --- | --- | --- | --- | --- | --- | --- |
|  |  | RNA | ATAC |  |  | RNA | ATAC |
| PBMC | Peripheral Blood | 9,378 | 9,378 | 1 | Yes | 14 | 14 |
| BMMC | Bone Marrow | 69,249 | 69,249 | 13 | Yes | 22 | 22 |
| Kuppe-2022 | Heart | 132,085 | 37,096 | 25 | No | 11 | 8 |
| Muto-2021 | Cortex of Kidney | 19,985 | 24,205 | 5 | No | 13 | 13 |
| Wilson-2022 | Cortex of Kidney | 22,944 | 44,995 | 8 | No | 13 | 16 |

We compared AtlasOT against state-of-the-art methods, including scGLUE^28^, a graph neural network (GNN) algorithm, which has consistently outperformed all other methods in three previous benchmarks^10,12,13^; and UINMF^29^, which was top-performing in a previous benchmark not including scGLUE^11^. We also included the closely related OT-based frameworks MOSCOT^18^ and scConfluence^20^. The batch correction algorithm Harmony^30^ is adopted as a baseline. None of these approaches allow for sample-specific mapping, so they were executed separately for each sample to satisfy the sample-specific matching constraint. We also evaluated two variants of AtlasOT. AtlasOT(UOT), which represents an unbalanced version of the OT only using the shared space, and AtlasOT(UGW), which only considers the unbalanced Gromov-Wasserstein using modality-specific spaces. We use the cell type annotation from both RNA and ATAC modality provided in the studies describing the respective data as the true labels. Methods were evaluated regarding their label-transfer accuracy.

We first leveraged paired multiome data (BMMC) and a single sample multiome dataset (PBMC), which are commonly used in benchmarking studies^11–13^, to optimize two main parameters of AtlasOT: the entropic regularization parameter *ε* of the efficient Sinkhorn’s optimization algorithm^31^ and the *α* controlling the trade-off between the GW and OT formulations (Supp. Fig. S1). These results showed no clear impact from *ε*, while values below 10^−6^ were prone to numerical instability. Regarding *α*, a value of 0.9, which balances both GW and OT formulations, was optimal for both label-transfer accuracy and modality-integration (FOSCTTM)^17^ values. These were adopted in our experiments.

When considering the combined performance in all three imbalanced datasets, AtlasOT achieved the highest median labeltransfer accuracy among the eight evaluated methods (Fig. 2a), reaching 0.85, followed by its unbalanced variant AtlasOT(UOT) (0.82) and by scGLUE (0.81). Compared with AtlasOT(UOT), scGLUE, UINMF, scConfluence, MOSCOT, AtlasOT(UGW) and Harmony, the advantage of AtlasOT corresponds to relative improvements of 3.78%, 5.26%, 20.94%, 28.48%, 34.09%, 396.85% and 967.47%, respectively. Notably, AtlasOT also outperformed its unbalanced variant AtlasOT(UOT), indicating that the combination of shared-space and modality-specific information is beneficial. Harmony performed the worst on this task, consistent with the fact that it was originally proposed as a batch-correction method on the RNA gene-expression matrix, rather than as a diagonal integration method. Examples of low-dimensional embeddings related to mappings are shown in Fig. 2c and Supp. Fig. S2.

**Figure 2.**
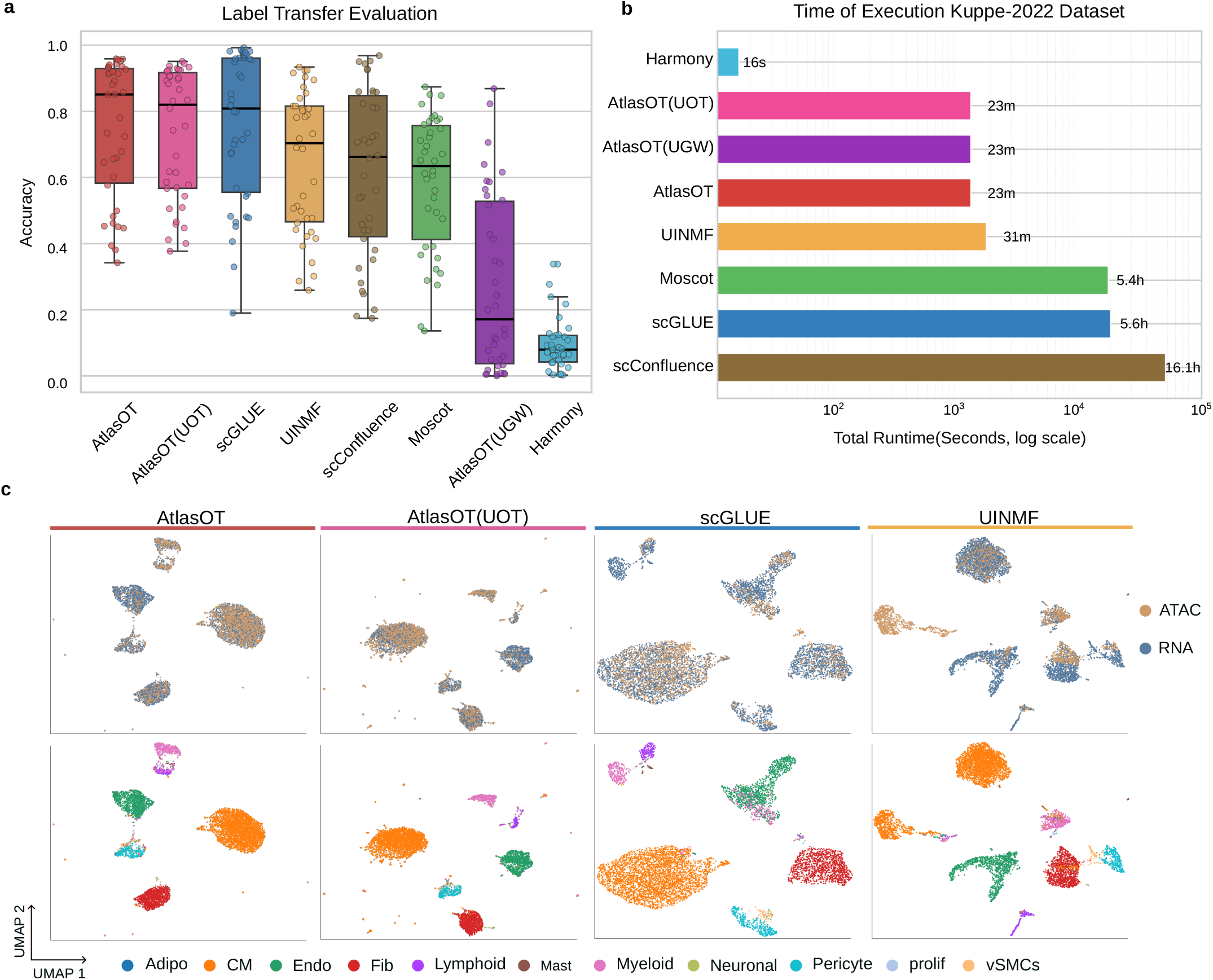
a) Boxplot with the RNA-to-ATAC label transfer accuracy (y-axis) for all evaluated datasets and algorithms. Methods are sorted by decreasing median accuracy. **b)** Runtime benchmarking of each method on the RNA-ATAC task using the Kuppe-2022 dataset. **c)** Example UMAPs with integrated datasets and cell type annotation for a selected sample RZ_P11 from Kuppe-2022 for the four top performing algorithms.

We additionally evaluated the runtime of each method on the RNA–ATAC task using the Kuppe-2022 dataset (Fig. 2b). Harmony was the fastest (0.3 min), closely followed by the three AtlasOT variants (23.1 min each), then UINMF (31.0 min), MOSCOT (321.4 min), scGLUE (336.5 min), and scConfluence (963.0 min). Notably, although Harmony was the fastest, it had the lowest label-transfer accuracy (Fig. 2a), whereas scConfluence, despite GPU acceleration, was the slowest method. Altogether, these results strongly support the value of AtlasOT in the accurate and efficient mapping of imbalanced, sample-matched scRNA-scATAC data.

### Label transfer allows detection of rare cells

It is common in Atlas data that modalities with lower cell counts have fewer cell types detected, possibly due to the difficulty in delineating rare cells. This is the case for our most imbalanced datasets. In the heart myocardial infarction data^4^, eleven cell types were detected in RNA-seq vs. eight cell types in ATAC-seq. For Wilson-2022, sixteen cell types were found in the more abundant ATAC-seq vs. thirteen in the RNA-seq. To address this, we evaluate the performance of AtlasOT in the detection of rare cells by using the OT mapping to perform label transfer. Next, we estimate and evaluate gene markers associated with the newly detected cells. Mast and adipocyte cells, which represented the lowest-frequency cells in the scRNA-seq data (0.51% and 0.26%, respectively), could not be detected in scATAC. We used label transfer from scRNA-seq to scATAC from AtlasOT to reassign cell types and checked ATAC-seq based gene activity scores on the new cell labels. We observe that new cell type labels reflect cell distributions (Fig. 3a), while the confusion matrix indicates that adipocytes were initially labelled as cardiomyocytes, whereas mast cells were labelled as myeloid cells. When considering marker genes, we observe clear mast cell markers (KIT, FCER1A) and adipocyte markers (PDK4, LPL) in the relabelled cell clusters (Fig. 3c). A similar analysis in the Kidney datasets (Wilson-2022) (Fig. S3 a-c) also indicates that label transfer can split the group of immune cells into B, T, and monocyte cells. Altogether, these results support the relevance of the mapping from AtlasOT.

**Figure 3.**
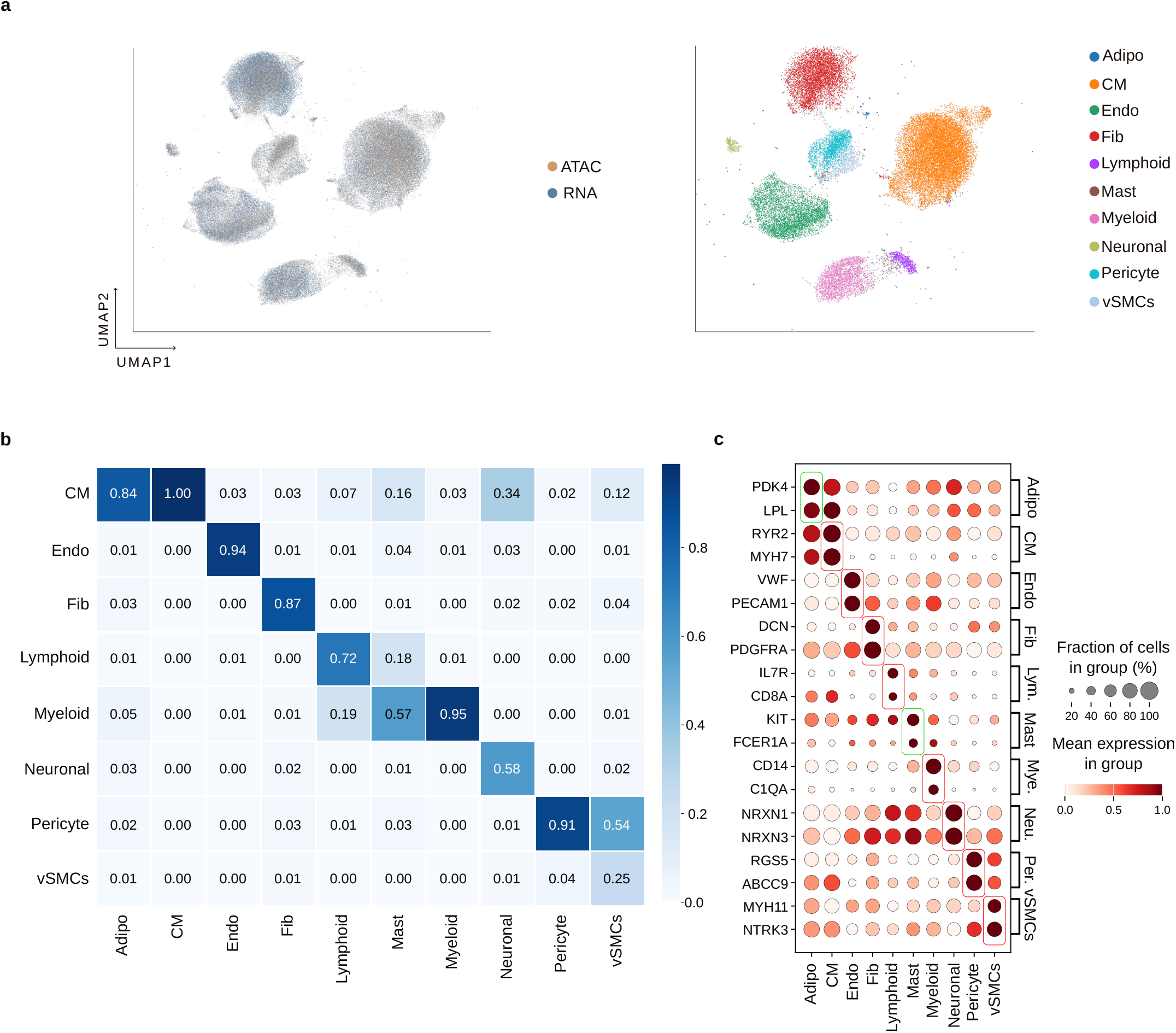
a) UMAP with all samples from the Kuppe-2022 heart data showing the two modalities (left) and cell types after transfer (right) as estimated by AtlasOT. **b)** The confusion matrix showing how cell labels change from the original (y-axis) vs. transferred (x-axis) cell types. **c)** Dot plot showing the gene activity of cell marker genes (y-axis) vs. scATAC-seq cell types after label transfer (x-axis). Adipocyte and mast cell markers are boxed in green, all others in red.

### Benchmarking of scRNA-ST mapping in disease atlas

We next evaluate AtlasOT and competing approaches in the problem of sample-specific mapping of single-cell data to spatial transcriptomics. AtlasOT formulates this task as a diagonal integration problem and uses the same FUGW objective function as in the RNA–ATAC integration, but introduces an additional feature tailored to the ST setting. For the ST-specific modality space, AtlasOT adopts a spatially constrained *k*NN graph, i.e., only adjacent edges are considered in the *k*NN graph. Moreover, for the imputation task, AtlasOT uses a smoothing function in the imputed expression matrix to enhance spatial continuity (see Methods). To benchmark AtlasOT and competing approaches, we adopted the feature imputation problem as usually performed in RNA-ST mapping studies^14^.

Methods were evaluated in two multimodal disease atlases with scRNA-seq and Visium-based spatial transcriptomics on myocardial infarction (Kuppe-2022^4^) and breast cancer (HTAPP^32^) with 26 and 15 samples, respectively (Table 2). Moreover, as expected, Visium experiments have sparser values (Table 2), with fewer average genes detected per spot than in the single-cell data. This supports the fact that one modality has a higher signal-to-noise ratio than the other.

**Table 2.**
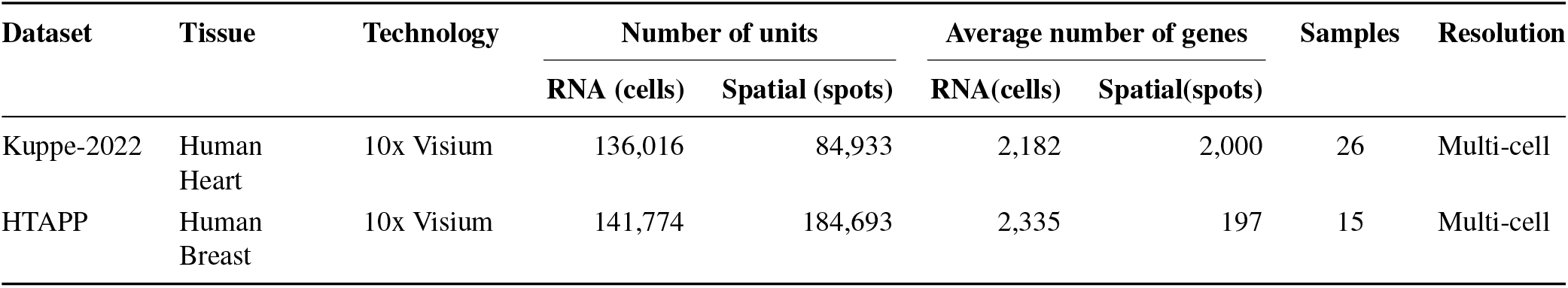
Summary of scRNA-seq and Spatial Transcriptomics datasets used for multimodal integration and spatial imputation.

These were contrasted with six SOTA methods spanning the mainstream algorithmic paradigms: the deep-learningbased algorithms Tangram^33^ and gimVI^34^, the optimal-transport-based NovoSpaRc^19^ (as implemented in MOSCOT^18^), scConfluence^20^ and SpaOTsc^35^, and the manifold-alignment-based SpaGE^36^. This selection includes the three top-performing approaches (Tangram, gimVI, and SpaGE) in a spatial mapping benchmarking^14^.

For the imputation evaluation, first, genes are held out from the ST dataset, and their expression is estimated after scRNA-seq-to-ST mapping. For this, we select the top 100 highly variable genes per dataset using Moran’s I and split these into 10 folds (the held-out test set). The imputation was evaluated with four complementary metrics: PCC (Pearson Correlation) and RMSE (Root Mean Square Error) measuring numerical agreement, JS (Jensen-Shannon divergence) measuring distributional similarity, and SSIM (Structural Similarity Index) similar to that in^14^.

We developed a parameter optimization approach using nested cross-validation on the training data to optimize AtlasOT parameters in a data-specific manner. In short, we selected genes not included in the test set and performed a parameter sweep on this validation gene set to identify the optimal parameters, which were applied across all samples (see Methods). Note that this unsupervised approach is implemented as a feature of AtlasOT and can be applied to any new incoming dataset.

The pooled results across the two datasets are summarized in Fig. 4. On PCC and RMSE, the two primary metrics measuring numerical agreement, AtlasOT achieves the highest mean among the seven methods (Fig. 4a,d), reaching average PCC = 0.203 and RMSE = 1.243, while the runner-up NovoSpaRc achieves 0.181 and 1.263, corresponding to relative improvements of 12.15% and 1.58%, respectively. In terms of the Jensen-Shannon metric, which measures the similarity between predicted and estimated expression values, AtlasOT ranks second behind Tangram (Fig. 4b). Finally, on the SSIM metric, which evaluates the structural similarity of the image, AtlasOT has the best performance, reaching an average SSIM = 0.148, while the runner-up SpaGE achieves 0.133, corresponding to a relative improvement of 11.54%. Overall, AtlasOT not only dominates three of the four complementary metrics as the top performer but also consistently maintains a leading position across all evaluations.

**Figure 4.**
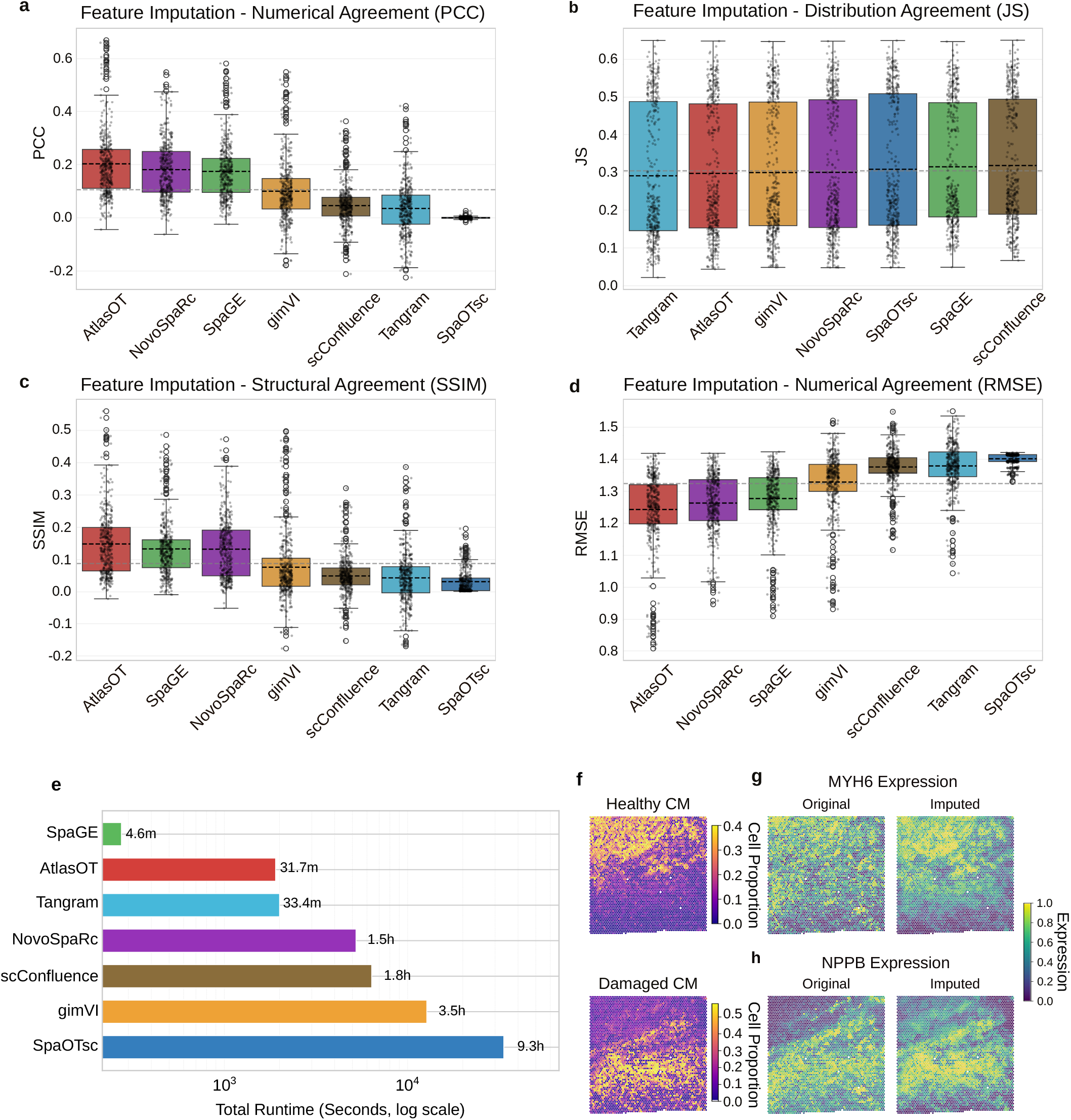
a) Boxplot with the Pearson Correlation Coefficient (PCC; y-axis) for all evaluated datasets and algorithms on the scRNA-ST gene imputation task. Methods are sorted by decreasing average PCC. **b)** Boxplot with the Jensen-Shannon divergence (JS; y-axis) for all evaluated datasets and algorithms. Methods are sorted by increasing average JS. **c)** Boxplot with the Structural Similarity Index (SSIM; y-axis) for all evaluated datasets and algorithms. Methods are sorted by decreasing average SSIM. **d)** Boxplot with the Root Mean Square Error (RMSE; y-axis) for all evaluated datasets and algorithms. Methods are sorted by increasing average RMSE. Altogether higher PCC and SSIM indicates best results, while lower JS and RMSE indicates best results. **e)** Runtime benchmarking of each method on the RNA-ST task using the Kuppe-2022 dataset. **f)** Deconvolution abundance of cardiomyocytes cell subtypes in a border zone sample. **g)** Expression values of the original and imputed MYH6 gene in a border zone sample. **h)** Expression values of the original and imputed NPPB gene in a border zone sample.

We further benchmarked the runtime on the RNA-ST task using the Kuppe-2022 dataset (Fig. 4e). SpaGE was the fastest (4.6 min), with AtlasOT ranking second (31.7 min), followed by Tangram (33.4 min), NovoSpaRc (87.1 min), scConfluence (105.9 min), gimVI (212.0 min), and SpaOTsc (557.6 min). Although SpaGE was the fastest, its imputation performance was markedly lower than AtlasOT on all four metrics (Fig. 4a–d). Together with its top performance in label transfer and gene imputation, these results show that AtlasOT achieves a favorable balance between computational efficiency and integration performance.

### Marker gene imputation in Spatial data

We next evaluated the performance of AtlasOT in leveraging spatial data to recover biologically relevant information in space. In the RNA-ST integration task, the mapping **T** from AtlasOT can be used for two tasks: it can project RNA gene expression onto the spatial dimension for imputation (Eq. 7) and it also represents the proportion of each cell type within each spot (or cell type deconvolution (Eq. 9)). As an example, we use Visium slides characteristic of ischaemic tissue and border zone tissue (the area between healthy and infarct tissues). In the border zone sample (Fig. 4f), we observe that cell deconvolution supports a clear transition from healthy CM to damaged CM regions. Moreover, imputation of relevant genes related to healthy (MYH6) and damaged myocardium (NPPB) also relates well to the cell deconvolution (Fig. 4g-h). In the ischaemic sample, deconvolution results distinguish the spatial locations of the progenitor-like SCARA5 fibroblasts and disease-transformed myofibroblasts (Supp. Fig. S4). Moreover, gene imputation shows that AtlasOT can improve the gene expression of the progenitor fibroblast marker SCARA5 (Supp. Fig. S4), which could not be detected in the original work introducing the data^4^, due to its sparsity.

### Spatial chromatin flow using the snRNA-snATAC and snRNA-ST mapping

To dissect regulatory mechanisms underlying tissue remodelling in disease atlases using multimodal single-cell data, understanding which transcription factors drive these gene expression patterns is key^37^. Chromatin potential also called chromatin velocity, which measures how chromatin accessibility changes precede expression changes of downstream genes, is crucial for such a task^38–41^. While these approaches were successfully applied in multimodal single-cell data, they require computationally derived pseudotime estimates and were rarely used in the spatial context due to the lack of spatial multimodal data. AtlasOT leverages the snATAC-to-snRNA and snRNA-to-ST mapping to estimate TF activity and TF regulon expression in space. These can be used to estimate chromatin flows corresponding to regions where chromatin changes precede regulon expression. Finally, we provide for each TF a chromatin flow score based on the proportion of the tissue covered by spatially coherent flows leading to basin regions (Fig. 5a; Methods Section).

**Figure 5.**
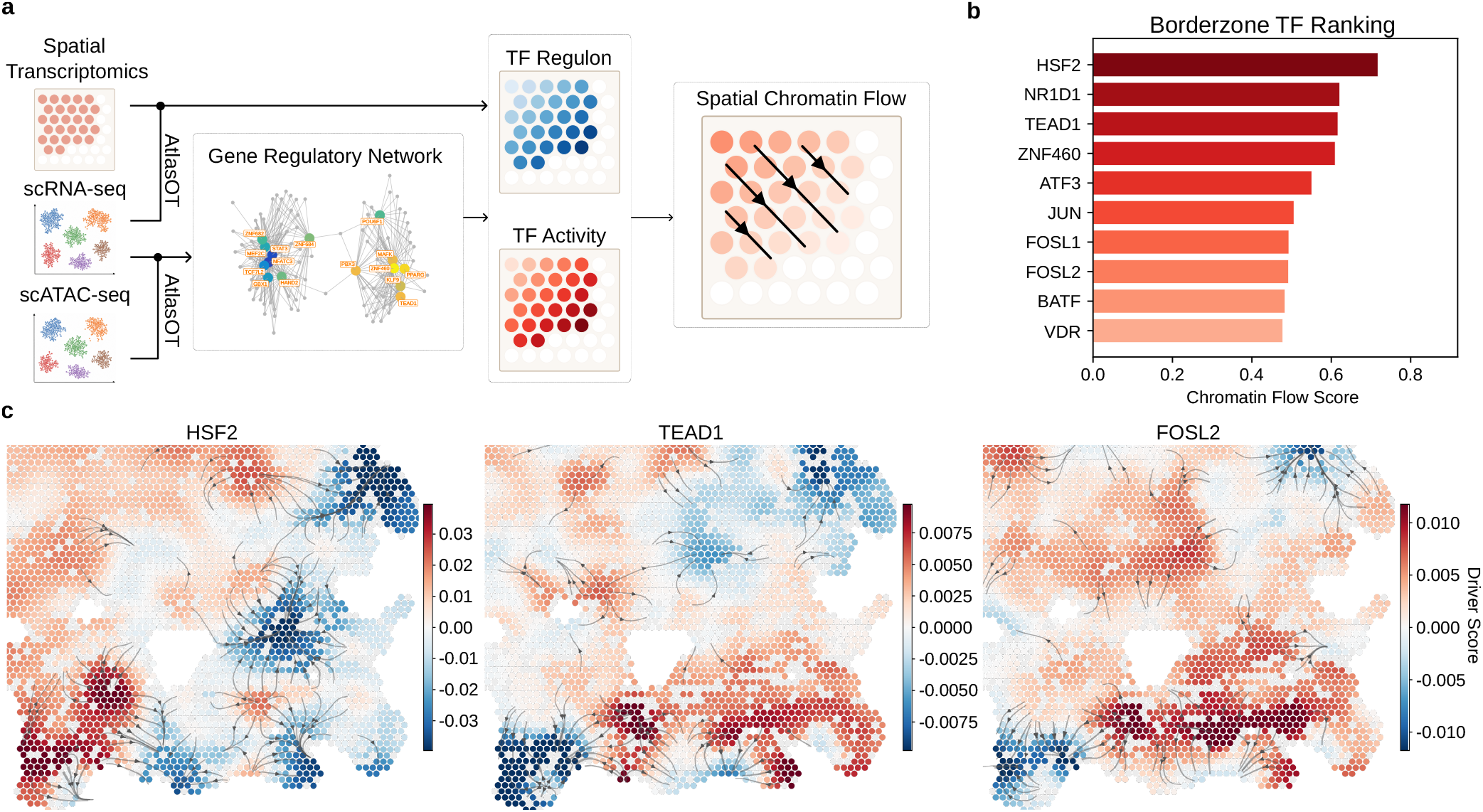
a) Schematic of the spatial chromatin flow workflow. AtlasOT maps snRNA-seq to snATAC-seq and, independently, snRNA-seq to ST within the same sample; chaining the two transport plans yields a sample specific snATAC-to-ST correspondence. First, AtlasOT creates an enhancer-based GRN to obtain TF regulons. This allows the TF activity and TF regulon expression to be mapped into the tissue coordinate system. Next, AtlasOT estimates chromatin flows by finding regions where TF activity precedes the TF’s regulon expression, and a chromatin flow score indicating TFs with highest spatially coherent flows. **b)** Bar plot showing the TFs with the ten highest chromatin flow scores. Here, we only consider TFs whose TF activity is positive with respect to their regulons (activators) and increases over pseudotime, i.e. associated with damaged cardiomyocytes. Significance is assessed by a spatial permutation test (Eq. 17). **c)** Spatial chromatin flows of selected top-ranked TFs: HSF2, TEAD1 and FOSL2.

As a case study, we focus on the border zone sample RZ_BZ_P3, as it represents a cardiomyocyte-rich area, where there is a clear remodelling of cardiomyocytes (CM) from healthy to disease states (Supp. Fig. S5). Next, we estimate the eGRN from the snRNA-snATAC maps (Supp. Fig. S5a) and only consider TFs with increased TF activity and TF regulons in damaged cardiomyocytes. Finally, we use our chromatin flow score to rank candidate TFs by the spatial extent of their coherent flow (Fig. 5b). The resulting chromatin flow fields of selected top-ranked TFs are shown in Fig. 5c. HSF2 is associated with heat-shock stress conditions and has been linked to hypertension-induced cardiac hypertrophy and heart failure in mouse models^42^. TEAD1 is part of the Hippo pathway and a known regulator of embryonic cardiomyocytes. Recent studies of TEAD1 indicate its overexpression leads to cardiomyocyte remodelling and cardiomyopathy^43^. Moreover, five of the top TFs belong to the AP-1/bZIP family, reflecting a coordinated cardiac stress-remodelling program, as supported by the literature indicating that JUN drives adaptive hypertrophy while FOSL2 mediates the maladaptive oxidative-stress and fibrotic arms^44,45^. The prominence of this family alongside the top three TFs suggests that the overall signature reflects a coordinated regulatory program anchored by HSF2 and TEAD1 as upstream or parallel regulators rather than a set of independently acting, disease-risk-associated factors.

## Discussion

Our study proposes AtlasOT, an optimal transport framework based on a unified Fused Unbalanced Gromov-Wasserstein mathematical formulation, designed for the integrated tasks of RNA-ATAC label transfer and RNA-ST gene imputation in disease atlases. AtlasOT simultaneously addresses three key limitations of existing methods: modality heterogeneity arising from distinct feature spaces, cell population imbalance inherent to real-world atlas data, and the need to constrain mapping to cells from the same sample or donor.

In the RNA-ATAC label transfer task, AtlasOT achieved the highest median label-transfer accuracy (0.85) across three disease atlases, outperforming all evaluated methods, including scGLUE, which has consistently ranked first in prior benchmarks. A recurrent challenge when performing multimodal label transfer is the differences in single-cell recovery in each of the data modalities. These differences were also observed across all evaluated datasets, where the modality with fewer recovered cells consistently displayed fewer detected cell types. In this regard, AtlasOT’s mapping recovered these missing rare populations and confirmed their identity via known marker genes. Furthermore, we note that all existing benchmarking studies and challenges for unpaired scRNA-scATAC integration rely on artificially balanced datasets, a consequence of using paired multiome data as a gold standard^10–13^. In contrast, our benchmark supports reliable evaluation by accounting for the imbalance found in real atlas studies.

In the RNA-ST imputation task, AtlasOT was the top performer on three of four complementary metrics and ranked second on Jensen-Shannon divergence. We further illustrated AtlasOT’s applicability through gene imputation and cell-type deconvolution on selected spatial samples, recovering spatial patterns for markers not resolved in the original study describing the data. Finally, unlike other methods that can only align modalities in a shared latent space, by chaining the RNA-ATAC and RNA-ST transport plans, AtlasOT enabled a spatial chromatin flow analysis that projected transcription factor motif activity and target gene expression onto a shared spatial coordinate system, revealing HSF2, TEAD1 and FOSL2 as TFs associated with the regulation of cardiac remodelling in a myocardial infarction border zone sample.

Altogether, AtlasOT provides a unified, biologically constrained framework for integrating sample-matched multiomics data in disease atlases, explicitly accounting for the imbalance and donor-matching constraints that characterize real-world data but are absent from most existing benchmarks. AtlasOT is also among the fastest methods on both tasks, requiring 23.1 min for RNA-ATAC and 31.7 min for RNA-ST, while ranking first or second on every evaluated metric. Future work will focus on extending validation to additional tissues and modalities such as histology or protein abundance, and further characterizing the computational scalability of the framework for population-scale disease atlases.

## Methods

### Notations

AtlasOT takes two modalities as input to be integrated: a source and a target, each of which can be represented as 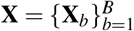 for the source modality and 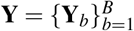 for the target modality. Here, 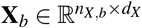 represents the cell-by-feature matrix of the source modality with donor *b* and 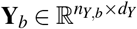 represents the cell-by-feature matrix of the target modality for donor *b*. We assume here that all modalities have cells from all samples/donors. Typical multimodal single-cell and spatial atlases pose three key challenges: (1) data in different feature spaces, i.e., *d*_*X*_ ≠ *d*_*Y*_; (2) unbalanced cell numbers, i.e., *n*_*X,b*_ ≠ *n*_*Y,b*_; (3) the mapping should be constrained to the same sample or donor. The main computational task is to find a mapping between cells across **X** and **Y**, such that only cells from the sample *b* are mapped. More formally, the coupling matrix 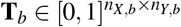 represents the optimal transport plan between the source and target data for the donor/sample *b*.

### AtlasOT

#### Cross-Modal Alignment with FUGW

Optimal Transport (OT)^25^ provides a powerful computational framework for aligning data distributions by defining the cost matrix between the source data and the target data, and then deriving an optimal transport solution to map the source distribution to the target distribution. However, in multimodal single-cell analysis, source and target data often reside in distinct feature spaces, making it infeasible to directly compute a transport cost matrix between them, unless one applies lossy transformations of the data. To address this, the Gromov-Wasserstein (GW)^24^ method was introduced, which aligns data by leveraging modality-specific metric spaces without requiring them to share a common feature space. While effective, GW neglects potential biological correspondences or explicit similarities across modalities. The Fused Unbalanced Gromov-Wasserstein (FUGW)^23^ framework integrates the strengths of both OT and GW. Moreover, the use of an unbalanced variant, which allows the masses of the source and target distributions to differ, offers additional flexibility due to cell capture differences in distinct omics modalities. For a sample index *b*, the two inputs are both modelled in a shared latent space. Finally, the *KL* is the Kullback-Leibler divergence, which is used to relax the distribution constraints, allowing for an unbalanced transport.

More formally, for a given sample *b* ∈ {1,…, *B*}, we have:

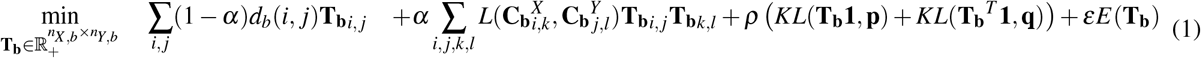

where *d*_*b*_(*i, j*) represents the cost matrix computed between a pair of observations *i* and *j* in the source and target data in the shared space *Z*, the transport plan **T**_**b**_ encodes the optimal mapping from source cells to target cells and the term 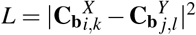 defines the loss function that quantifies the discrepancy between the intra-modality similarity structures, where *i, k* index cells in the space of the source cells *X* and *j, l* index cells in the space of the target data *Y*. **C**_**b**_^*X*^ and **C**_**b**_^*Y*^ are modality-specific spaces defined below. An entropy regularization term *E* is incorporated to improve computational efficiency during optimization^31^. **p, q** are uniform distributions representing a weak prior on the source and target distributions.

The hyperparameter *α* ∈ [0, 1] controls the importance of the optimal transport vs. the Gromov-Wasserstein terms. For *α* = 0, we obtain a standard OT only considering the shared space; and when *α* = 1 we have a GW, which only considers the modality specific spaces. The hyperparameter *ρ* ∈ [0, +∞) is the relative importance of the distribution constraints, i.e. lower values allow a higher unbalance between the mappings. *ε* ∈ [0, ∞) controls the relative importance of the entropy loss, where high values tend to the balanced OT formalization.

#### Cross-modality shared space

The OT component (left term of Eq. 1) requires the definition of a distance *d*_*b*_(*i, j*) in a shared space *Z* representation. This is not always trivial to estimate in cases where feature descriptors of **X** and **Y** are distinct, i.e., scRNA/ST have genes as features and scATAC-seq has genomic peaks as features. A surrogate for gene expression in scATAC-seq is the gene activity score^46^, which quantifies the accessibility of peaks around a gene. This measurement is lossy, i.e., only partially reflects gene expression, due to the lower sequencing depth of scATAC and the difficulty in assigning peaks to genes. Similarly, for ST, the number of genes measured is usually lower than in scRNA-seq (Table 2), which indicates higher noise levels in ST experiments. Therefore, we devise the following procedure to build a shared space by focusing on the more discriminative scRNA-seq modality. Moreover, AtlasOT does not consider the donor identity of cells for estimation of the shared space.

##### scRNA-scATAC shared space

Given the complete input matrix 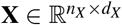 and target matrix 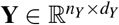, we first identify the intersection of their gene sets to obtain 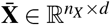 and 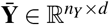, where *d* min(*d*_*X*_, *d*_*Y*_).

Given that scRNA-seq data 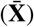 typically exhibits lower noise levels, we perform PCA on 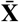 and retain the top *k* principal components where *k* is determined by retaining components with explained variance ratio above 10^−3^. Let **V**_*k*_ ∈ R^*d*×*k*^ be the loading matrix of the top *k* components.

Next, we project both source 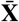 and target 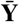 into this shared PCA space.

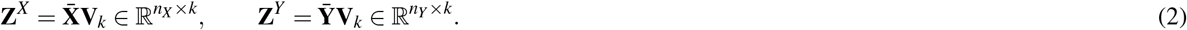

Of note, this operation is done ignoring sample labels. Subsequently, for each donor *b*, we extract the per-sample embeddings 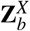 and 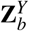 from 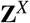 and 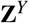 and compute the cross-modality cost matrix **d**_*b*_(*i, j*) (used in the OT term of FUGW) by computing the cosine distance between the shared-space embeddings:

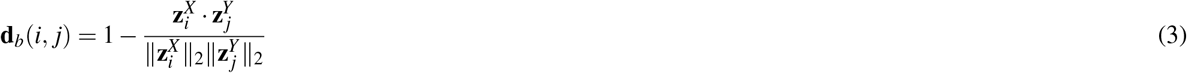

where 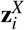 and 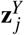 are rows of 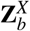 and 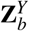, respectively.

##### scRNA-ST shared space

The shared space for scRNA-ST is constructed similarly: PCA is performed on the scRNA-seq data, and both scRNA-seq and ST data are projected onto the same principal component space. While scRNA-seq and ST are likely to be in the same feature space (genes), the step of finding common genes is still needed, as the gene number might differ due to platform differences or the adoption of gene panels in the ST modality.

#### Modality specific spaces

To capture the manifold structures inherent in single-cell data and provide topological constraints for Gromov-Wasserstein alignment, we construct intra-modality geodesic distance matrices 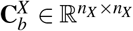 and 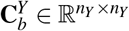 for each sample separately, inspired by ISOMAP. We first obtain modality-specific low-dimensional representations 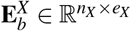 and 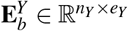. For scRNA-seq and scATAC-seq, data is reduced via PCA and an NMF algorithm^47^, respectively.

For scRNA-seq and scATAC-seq, we construct a *k*-nearest neighbour (*k*NN^48^) graph *G* (*k*_*nn*_ = 30) directly in feature space. The edge weights are defined by the cosine distance 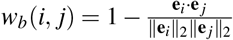, where **e**_*i*_, **e** _*j*_ are rows of 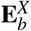 (or 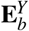, respectively).

For spatial transcriptomics data, unlike dissociated single-cell data where only expression similarity matters, spatial spots that are physically adjacent may not share similar expression profiles (e.g., across tissue boundaries), while spots with similar expression may be far apart. We therefore incorporate physical constraints. Let **S**_*b*_ ∈ R^*n*×2^ represent the spatial coordinates of *n* spots for sample *b*. We first define graph adjacency by constructing a *k*-nearest neighbour graph (*k*_phys_ = 15) based on Euclidean distances between spots in **S**_*b*_, thereby restricting edges to physically proximate spots. For any connected edge (*i, j*), the edge weight is then assigned using the cosine distance between the feature vectors **e**_*i*_, **e** _*j*_ (rows of **E**_*b*_).

Next, for all modalities, we use Dijkstra’s algorithm^49^ to compute all-pairs shortest paths on the respective graphs, yielding the intra-modality geodesic distance matrices 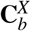 and 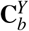 used in the FUGW objective (Eq. 1). Due to the presence of isolated data “islands” during *k*NN graph construction, the resulting geodesic distances are essentially local. To prevent infinite distances caused by disconnected components, we replace ∞ with 1.5 times the maximum detected distance. We define the final cost matrix as:

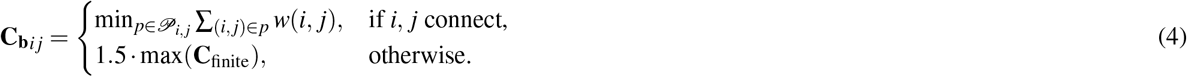

This construction is applied independently to each modality, yielding 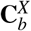 and 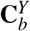 used in Eq. 1.

#### Computational Complexity

**AtlasOT** solves each sample independently. For a single sample with *n* source and *m* target cells, each FUGW iteration has complexity *O*(*n*^2^*m* + *nm*^2^), dominated by the matrix product 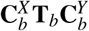 in the Gromov-Wasserstein gradient^23,25^.

Let 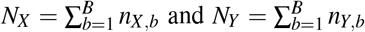 be the total number of source and target cells across all samples. Pooling all samples into a single transport problem would give 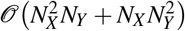, which is prohibitive for large atlases. By solving per-sample problems, AtlasOT achieves total complexity 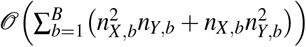. Since 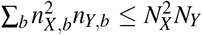, the per-sample strategy is always cheaper. Moreover, the per-sample solves can be parallelized providing additional speed ups.

#### Label Transfer and Joint Embedding

For certain modalities where label annotation is challenging due to signal sparsity, we leverage the transport plan **T** to transfer labels from an alternative modality. For notation simplicity, we drop the sample specific term *b* hereafter.

Given the dimensionality reduction matrices for the source modality 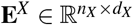 and the target modality 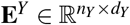, we obtain the transport plan 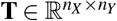 via AtlasOT. To transfer a specific label from the source to the target modality, the label is first converted into a one-hot encoded vector 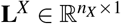. Since **T** represents the transition probabilities between each source point and each target point, the label transfer results can be computed directly. This process is likewise applicable in the reverse direction. The specific calculation formulas are as follows:

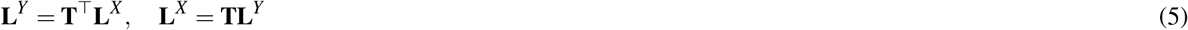

The transport plan is also applicable for joint embedding to support additional downstream analyses. This can be obtained

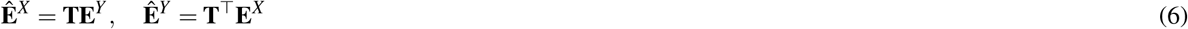

#### Spatial Gene Imputation and Deconvolution

We can also leverage **T** to impute genes; for example, we can use a scRNA-seq gene expression matrix to impute an ST gene expression matrix. Specifically, for a given scRNA-seq modality of a sample 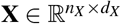, the formula for imputing the ST gene expression matrix Ŷ is as follows:

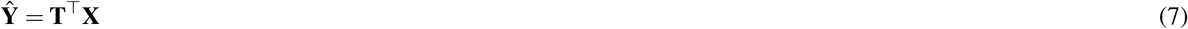

We next use random walk smoothing on the imputed gene expression to obtain smoothed estimates. For this, we use the spatially constrained graph (see above) using the cosine distance of spot features as edge weights. Then we use the cosine distance *w*(*i, j*) to construct an affinity matrix **A**_*i, j*_ = *e*^−*w*(*i, j*)^ between nodes. We then obtain the degree matrix of **A, D**_*i, j*_ = Degree(**A**_**i**,**j**_). At the *t*-th iteration, the smoothing of **Ŷ** is performed as follows:

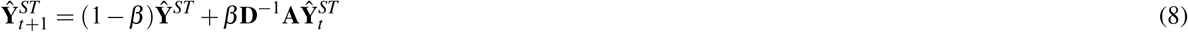

where *β* controls the weight of smoothing.

Furthermore, **T** can directly perform cell-type deconvolution. Let 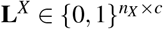 be a one-hot encoding of the cell-type labels for the scRNA-seq cells, where *c* is the number of cell types. The deconvolution matrix 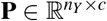 is obtained by projecting the labels through the transport plan:

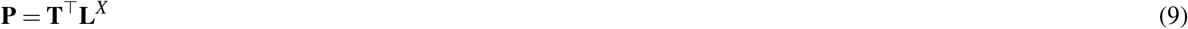

Each row of **P** reflects the estimated cell-type proportions within the corresponding spatial spot.

#### Spatial chromatin flow

AtlasOT leverages the mappings between RNA-ATAC and RNA-ST modalities provided by **T**_RNA_→_ATAC_ and **T**_RNA_→_ST_ to estimate TF activity and TF regulon scores. For TF regulon scores, we ran scMEGA^50^ on the AtlasOT paired RNA-ATAC cells to infer an enhancer-based gene regulatory network. For each transcription factor (TF), this provides us with the underlying regulons (target genes). For this, we use the RNA-ST mapping to estimate the imputed gene expression as described above (Eq. (7) and smoothed via Eq. (8)). The TF regulon score (**Ŷ** ^Regulon^) is the weighted average of all target genes, using the scMEGA TF-gene correlations as weights.

For the TF activity score, we first compute the transcription factor motif activity scores 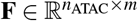 from scATAC-seq peaks using chromVAR^51^ with the JASPAR2020^52^ vertebrate motif database. Next, we leverage **T**_RNA→ATAC_ and **T**_RNA→ST_ to obtain a mapping between ATAC and ST, i.e.

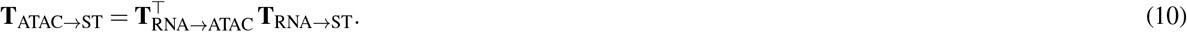

We can then use **F** and the imputation procedure (Eq. (7)) to estimate TF activity scores **Ŷ** ^TF^:

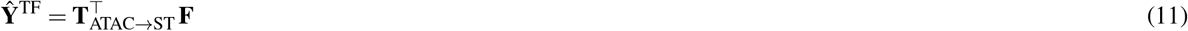

Finally, we performed rank-based normalisation of both the imputed regulon gene expression and the TF motif activity to the interval [0, 1], where 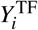 and 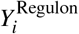 are rank-normalised TF activity and regulon scores for spot *i*. Moreover, as the eGRN is only specific to a cell type, we use the per-spot proportions from AtlasOT deconvolution (Eq. (9)) to make a target cell score 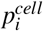. Within each spot, we then compute a smooth chromatin flow between the TF and its target gene regulon. As a starting point, we define

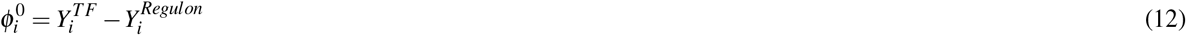

Next, we obtain spatially constrained smooth estimates as:

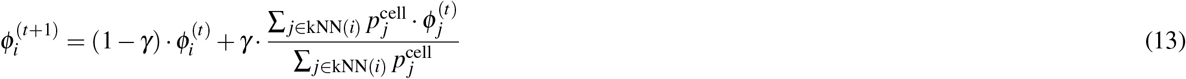

where *γ* = 0.6 and *kNN*(*i*) are the *k* nearest neighbours of *i*. By default *k* = 6 (hexagonal Visium lattice)and the algorithm uses 12 iterations. We then computed the chromatin flow **v**_*i*_ = −∇*ϕ*_*i*_ as:

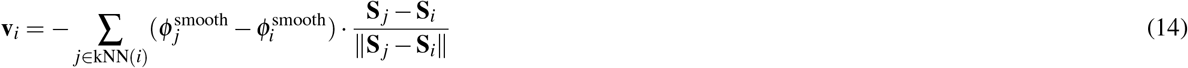

where **S**_*i*_ denotes the 2D spatial coordinate of spot *i*. As before^38,39^, we only consider spots where the TF motif activity exceeds the regulon gene expression rank 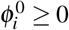.

We are interested in characterizing TFs whose coherent flow convergence covers the largest area of the tissue, namely basins. For this, we first identify the steepest-descent map *π* on the spatial *k*NN graph: each spot *i* is sent to whichever point in its closed neighbourhood {*i*}∪ *kNN*(*i*) has the smallest value of *ϕ*^smooth^.

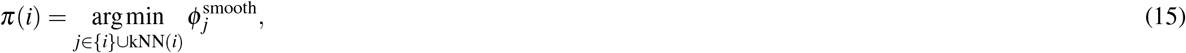

Iterating *π* from any spot produces a non-increasing sequence of *ϕ* ^smooth^ values on a finite domain, which must therefore converge to a fixed point or a spot with no lower-valued neighbour, i.e., a discrete local minimum of *ϕ* ^smooth^. Assigning every spot to the local minimum reached by iterating *π* partitions the target cell domain *D* into basins of attraction of the descent map.

Because *ϕ* ^smooth^ is estimated from a noisy spatial transcriptome, this partition typically contains many local minima that reflect small-amplitude fluctuations rather than genuine regulatory sinks. We therefore merge any minimum whose persistence, defined as the height difference between its basin bottom and the lowest saddle separating it from a neighbouring basin, falls below 5% of the dynamic range of *ϕ* ^smooth^ over *D*. The spots draining into the same surviving minimum *m* form a basin *B*_*m*_.

Finally, the chromatin flow score (CFS) quantifies the fraction of the tissue whose gradient field **v**_*i*_ drains into sinks.

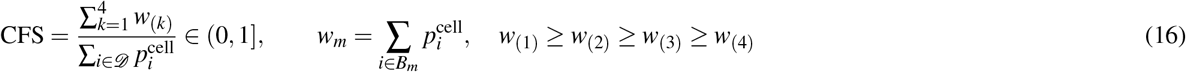

Here, a value near 1 indicates that nearly the entire cell-weighted domain converges, under steepest descent, to the m (4 as default) largest minima of *ϕ*^smooth^, representing a small number of spatially coherent regulatory fronts.

The significance of each CFS is assessed by a spatial permutation test (*N*_perm_ = 200): *ϕ* ^0^ is randomly permuted across spots, the full basin pipeline is re-run, and the empirical *p*-value is

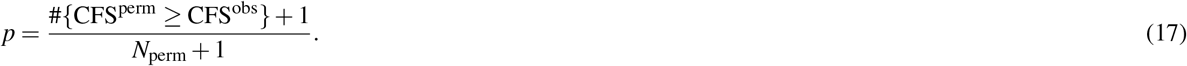

For visualization purposes, the filtered chromatin flow field is visualized as continuous streamlines in a single semitransparent dark grey colour following the scVelo^53^ convention, with line width proportional to flow speed and arrowheads indicating the direction of chromatin flow, from TF-driven regions towards the basin.

### AtlasOT hyperparameter selection

AtlasOT involves three hyperparameters: the fusion weight *α*, the unbalanced penalty *ρ*, and the entropic regularization *ε* (Eq. 1). For the RNA-ATAC task, *α* and *ε* were selected on the two paired multiome validation datasets (PBMC and BMMC), for which the true cross-modal cell correspondences are known, and then applied unchanged to the three unpaired test atlases (Kuppe-2022, Muto-2021, Wilson-2022). All FOSCTTM results reported below are averaged over the RNA-to-ATAC and ATAC-to-RNA directions. Due to the balanced nature of these data, this did not allow us to select the penalty value *ρ*, which was simply used based on the default value of 1.1.

To select *ε*, we monitored the total transported mass ∑_*ij*_ **T**_**b***ij*_ of the unbalanced coupling as a numerical diagnostic (Supp. Fig. S1a). Because mass cannot be created, the total mass must never exceed one; values below one are permitted and reflect the unbalanced relaxation. Smaller *ε* yields a sharper, more faithful coupling, but below *ε* = 10^−6^ the solver becomes numerically unstable, and the total mass exceeds one. We therefore set *ε* = 10^−6^, the smallest value at which the solver remains stable.

To select *α*, we scanned *α* ∈ [0, 1] on the validation datasets and quantified the resulting couplings with FOSCTTM (Eq. 19), which requires known ground-truth matches and is lower for better alignment (Supp. Fig. S1b). FOSCTTM was consistently minimized at *α* = 0.9, whereas *α* = 1 (pure GW) collapsed the alignment. We therefore fixed *α* = 0.9 for all test datasets.

For the RNA-ST task, we used the top 100 Moran’s-I-ranked genes as a held-out 10-fold test set; *α* and *ε* were tuned on the training genes and then evaluated on the held-out test genes. Specifically, we selected the next 50 ranked genes (ranks 101–150), disjoint from the test set, held out 20 of them as validation genes, and tuned the hyperparameters on three representative samples per dataset, yielding *α* = 0.6, *ε* = 10^−1^ for Kuppe-2022 and *α* = 0.2, *ε* = 10^−2^ for HTAPP.

### Baseline Methods

**Moscot**^18^ is an OT-based multiomics alignment method using the Fused Gromov-Wasserstein framework. For the shared space, Moscot employs standard scVI embeddings with default parameters to compute cross-modal distances between scRNA-seq and scATAC gene activity. This algorithm is used for the RNA-ATAC integration task. We followed the parameters and configuration from the official Moscot tutorial https://moscot.readthedocs.io/en/latest/notebooks/tutorials/600_tutorial_translation.html. Although Moscot also has a module for RNA-ST integration, it is just a reimplementation of the NovoSpaRc algorithm (see below) and was therefore not evaluated here.

**scGLUE**^28^ is a graph neural network (GNN) based multiomics integration method that learns a shared embedding via a variational autoencoder with a prior knowledge graph of regulatory interactions (linking ATAC peaks to nearby genes based on genomic proximity) to bridge the two modalities. scGLUE has consistently ranked first in multiple recent benchmarks^10,12,13^. scGLUE is only able to integrate RNA-ATAC modalities. We followed the parameters and configuration from the official scGLUE tutorial https://scglue.readthedocs.io/zh-cn/latest/tutorials.html.

**Harmony**^30^ is a batch-effect correction method, originally proposed for scRNA-seq data integration, and is adopted here as a baseline for RNA-ATAC cross-modal alignment. We ran Harmony (harmonypy v0.2.0) with its default parameters https://github.com/slowkow/harmonypy.

**UINMF**^29^ is a non-negative matrix factorization method for unpaired multimodal integration, extending the iNMF framework to cross-modal data, and ranked first in a benchmark^11^. UINMF is able to integrate RNA-ATAC modalities. We followed the parameters and configuration from the official UINMF documentation https://welch-lab.github.io/liger/articles/SNAREseq_walkthrough.html.

**scConfluence**^20^ combines autoencoder-based dimensionality reduction with unbalanced OT loss function for cross-modal alignment. scConfluence is able to integrate both RNA-ATAC integration task and the RNA-ST spatial mapping task. We followed the parameters and configuration from the official scConfluence tutorial https://scconfluence.readthedocs.io/en/latest/tutorials/RNA_ATAC_pbmc_tutorial.html and https://scconfluence.readthedocs.io/en/latest/tutorials/RNA_FISH_tutorial.html.

**SpaOTsc**^35^ is an OT-based method for inferring spatial gene expression patterns and is able to integrate RNA-ST modalities.

SpaOTsc formulates spatial alignment as a structured OT problem matching both expression and spatial distance matrices, but it lacks the fused *α*-interpolation between shared-space OT and modality-specific GW, as well as unbalanced relaxation and per-sample constraints present in AtlasOT. We followed the parameters and configuration from https://github.com/zcang/SpaOTsc/blob/master/short_tutorial/spaotsc_tutorial_short.ipynb.

**SpaGE**^36^ is a manifold-alignment-based method that links scRNA-seq and spatial transcriptomics data via *k*-NN regression, and ranked among the top three in a previous spatial benchmark^14^. We followed the parameters and configuration from its tutorial https://github.com/tabdelaal/SpaGE except for *n*_*pv*, as the default value led to numerical instability. The original SpaGE uses a fixed cosine similarity threshold of 0.3 to filter principal vectors (PVs). When the number of shared genes is small and the number of PVs is low, the similarity of all PVs may fall below this threshold, resulting in an effective PV count of zero and the algorithm failing to run. We made this threshold a tunable parameter and automatically reduced it to 0.1 when the number of PVs is less than 100, to ensure the robustness of the algorithm on small-scale feature sets.

**NovoSpaRc**^19^ is a Fused Gromov-Wasserstein OT-based spatial reconstruction method. NovoSpaRc participates in the RNA-ST spatial mapping task. NovoSpaRc relies on balanced FGW with strict mass conservation and constructs spatial topology based on pure physical coordinates, which easily causes mapping ambiguity at tissue boundaries. We followed the parameters and configuration from the official NovoSpaRc tutorial https://github.com/rajewsky-lab/novosparc/blob/master/reconstruct_drosophila_embryo_tutorial.ipynb.

**Tangram**^33^ is a deep-learning-based mapping method that maps scRNA-seq cells to spatial locations via a probabilistic scoring function. Tangram ranked among the top three in a previous spatial benchmark^14^. We followed the parameters and configuration from the official Tangram tutorial https://tangram-sc.readthedocs.io/en/latest/getting_started.html.

**gimVI**^34^ is a deep-learning based model for gene expression imputation, and ranked among the top three in a previous spatial benchmark^14^. gimVI is only able to map RNA-ST data. We followed the parameters and configuration from the official gimVI tutorial https://docs.scvi-tools.org/en/1.0.0/tutorials/notebooks/gimvi_tutorial.html.

### Datasets and Preprocessing

We curated six diverse benchmark datasets from recent single-cell multiomics studies: five datasets with multimodal single-cell transcriptomic (scRNA-seq) and chromatin accessibility (scATAC-seq) data and one additional dataset HTAPP with the scRNA-seq and spatial transcriptomic (ST), with the Kuppe-2022 dataset utilized in both benchmark tasks. Detailed dataset information is provided in Table 1 and Table 2.

We use all samples available in the original datasets. As an exception, for Kuppe-2022^4^, sample IZ_P3 was excluded from all analyses, as it contained only 91 cells with a median of 533 genes and 589 total counts per cell, all the lowest among the samples, indicating a failed cell capture. Sample FZ_P18 was excluded from the RNA–ATAC integration due to the fact that it mostly recovers cardiomyocyte cells (72.3% of its ATAC population vs. an average of 38.41% CM cells). Because quality-based exclusions are task-specific (sample IZ_P3 was removed from all analyses, and sample FZ_P18 from the RNA-ATAC integration only), the number of samples used per benchmark differs between tasks (Tables 1 and 2). All datasets were derived from human atlas samples and cover both balanced and imbalanced data types, providing a rich resource for cross-omics integrative analysis. Due to the heterogeneity of different modalities—such as sequencing depth, technical noise, and feature space differences—which may interfere with subsequent joint modelling, we applied a unified and standardized preprocessing pipeline to ensure comparability and robustness of the integration results across modalities. The key steps are as follows:

For single-cell RNA sequencing (scRNA-seq) data, we employed a preprocessing pipeline based on the Scanpy^54^ framework. First, we excluded mitochondrial genes to mitigate the impact of potential technical noise. We then normalised the raw expression matrices and performed a log1p transformation. Subsequently, we performed PCA on the processed data, retaining principal components with an explained variance ratio greater than 10^−3^, followed by L2 normalization. For RNA-ATAC integration tasks, we considered all the cells and used Harmony^30^ for sample-level batch-effect correction.

For scATAC peak matrices, peaks detected in fewer than 10 cells were removed using muon^55^. Then, the peak counts were normalised and log1p-transformed to stabilize the data distribution and mitigate the influence of extreme values. For the dimensionality reduction of ATAC samples, we utilized the scOpen^47^ function. We applied the Harmony function for sample-level batch correction. For scATAC gene activity matrices, we similarly applied total count normalisation followed by a log1p transformation to alleviate sparsity and enhance numerical stability.

For RNA-ST integration tasks, as the goal was to impute gene expression matrices, we filtered out scRNA-seq cells with fewer than 200 expressed genes to reduce noise. Additionally, because different tissues contain distinct spatial-specific signals, we processed the corresponding RNA sub-samples independently and did not use Harmony. For spatial transcriptomics (ST) data, we applied normalisation and a log1p transformation to the expression matrices. This was followed by PCA, where principal components with an explained variance ratio greater than 10^−3^ were retained and L2-normalised. This pipeline was executed independently per sample.

### Evaluation metrics

#### Accuracy

After deriving the transferred labels via Equation 5, we utilized the pre-annotated labels as the ground truth to evaluate prediction accuracy for the specific sample *b* with *n* cells/spots. The average accuracy is defined as:

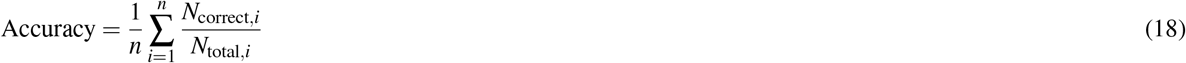

#### FOSCTTM

The Fraction of Samples Closer than True Match (FOSCTTM)^17^ evaluates the alignment quality on paired data with known cross-modal correspondences. For each source cell *i*, let 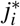 denote its true matching cell in the target modality. FOSCTTM is defined as:

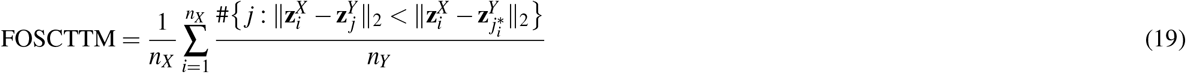

where #{·} counts the number of target cells *j* that lie strictly closer to cell *i* than its true match 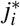, with distances measured by the Euclidean norm in the aligned embedding space. A lower FOSCTTM indicates better alignment and FOSCTTM = 0 corresponds to a perfect match in which every cell’s true match is its nearest neighbour.

#### PCC

After deriving the imputed gene expression vector **ĝ** ∈ ℝ^1×*N*^ via Equation 8, we evaluate its concordance with the ground truth vector **g** ∈ ℝ^1×*N*^ by computing the Pearson correlation coefficient (PCC):

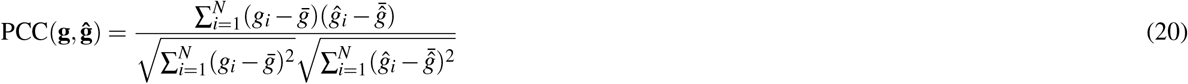

#### JS

The Jensen-Shannon (JS) divergence was employed to measure the distributional similarity between the imputed and ground truth gene expression profiles. A lower JS value indicates that the inferred expression proportions across all spots more closely resemble the true spatial distribution.

Since the JS operates on probability distributions, we first normalize each gene’s expression vector by its total sum across all spots:

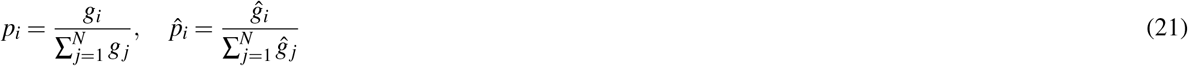

The JS is then calculated as follows:

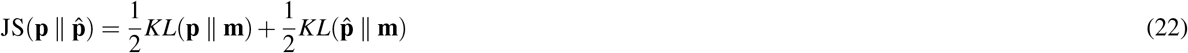

where 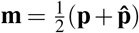 represents the pointwise mixture distribution, and *KL* denotes the Kullback-Leibler divergence.

#### SSIM

Structural Similarity Index Measure (SSIM) quantifies the structural fidelity between the imputed and ground truth spatial expression profiles. A higher SSIM score indicates better preservation of overall spatial patterns and boundaries.

Before calculating SSIM, we normalize the expression vector of each gene by its maximum value to scale the values into the [0, 1] interval:

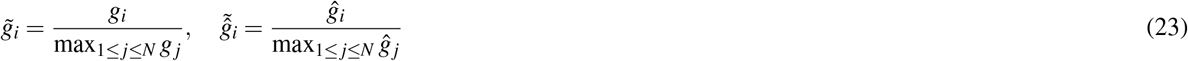

For a specific gene, the global 1D SSIM between its normalised ground truth vector 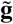 and imputed vector 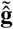 is computed as:

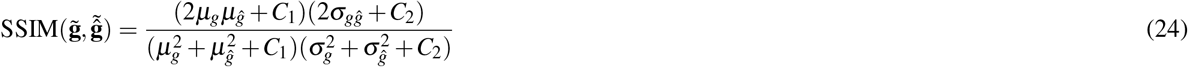

where *μ*_*g*_ and *μ*_*ĝ*_ denote the global mean values across all spots, 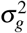 and 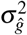 are their respective global variances, and *σ*_*gĝ*_ denotes their global covariance. The stability constants are defined as *C*_1_ = (*K*_1_*L*)^>2^ and *C*_2_ = (*K*_2_*L*)^2^, where *K*_1_ = 0.01, *K*_2_ = 0.03, and *L* = 1 is the dynamic range of the normalised expression values.

#### RMSE

Root Mean Square Error (RMSE) is a classic metric used to measure the absolute numerical difference between the imputed genes and the ground truth genes. A lower RMSE indicates that the absolute magnitude of the imputed expression is also very accurate. We first calculate the z-score for each imputed gene and ground truth gene, and then calculate the RMSE using the following formula:

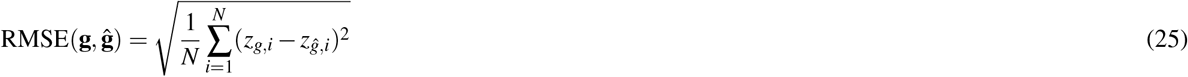

where *z*_*g,i*_ and *z*_*ĝ,i*_ represent the z-scored values of the ground truth and imputed expressions at the *i*-th spot, computed as 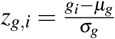 and 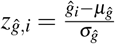, respectively (with *μ* and *σ* denoting the mean and standard deviation of the respective gene vectors across all *N* spots).

### Runtime Benchmarking Protocol and Computing Environment

We used the Kuppe-2022 dataset to measure the runtime of each algorithm. All runtime benchmarks were conducted on cluster compute nodes. For Harmony, SpaGE, NovoSpaRc, and SpaOTsc, experiments were run on a CPU compute node equipped with AMD EPYC 7502 32-core processors. For AtlasOT, scGLUE, scConfluence, Moscot, Tangram, and gimVI, we used a GPU compute node with one NVIDIA A100 40 GB GPU, with the remaining configuration identical to the CPU node. To ensure a fair comparison, all methods were uniformly restricted to at most 16 threads.

For the RNA-ATAC task, data preprocessing and dimensionality reduction are common steps for all algorithms and were therefore excluded from the total time. We measured only the time from invoking the algorithm to obtaining the integration result. For UINMF, which has special input data handling requirements, its full pipeline time was counted. In addition, AtlasOT requires extra computation of the shared space; this time, together with the subsequent AtlasOT runtime, is counted as the total time of AtlasOT.

For the RNA-ST task, all methods received identical preprocessed AnnData objects derived from the same upstream preprocessing pipeline. Timing covered the full process from benchmark instantiation and method invocation to the output of the imputed gene expression matrix (including any additional data preparation steps performed internally by each method). The computational cost of evaluation metric calculation is negligible.

## Data availability

The processed single-cell RNA, ATAC and spatial transcriptomics datasets generated and analysed in this study have been deposited in the Zenodo database under accession code https://zenodo.org/records/20713604.

## Code availability

AtlasOT is available as a Python package at https://github.com/CostaLab/AtlasOT

## Acknowledgements

This project has been funded by the German Research Foundation (DFG) (project GE 2811/3-2) and the Graphs4Patients consortia funded by the German Ministry of Education and Science (BMBF).

## Author information

K.P. and I.C. conceived the experiment(s), K.P. conducted the experiment(s), K.P., C.K. and I.C. analysed the results. J.N., M.R and B.C. supported software and method developments. All authors reviewed the manuscript.

## Competing Interests

The authors declare no competing interests.

## Footnotes

1 While OT defines one distribution as the source and other as the target, their interchange has no effect in the results.

## Notes

https://github.com/CostaLab/AtlasOT

### Competing Interest Statement

The authors have declared no competing interest.

