## Supplementary File for "AtlasOT - The Fused Unbalanced Gromov-Wasserstein for Multimodal Integration of Disease Atlases"

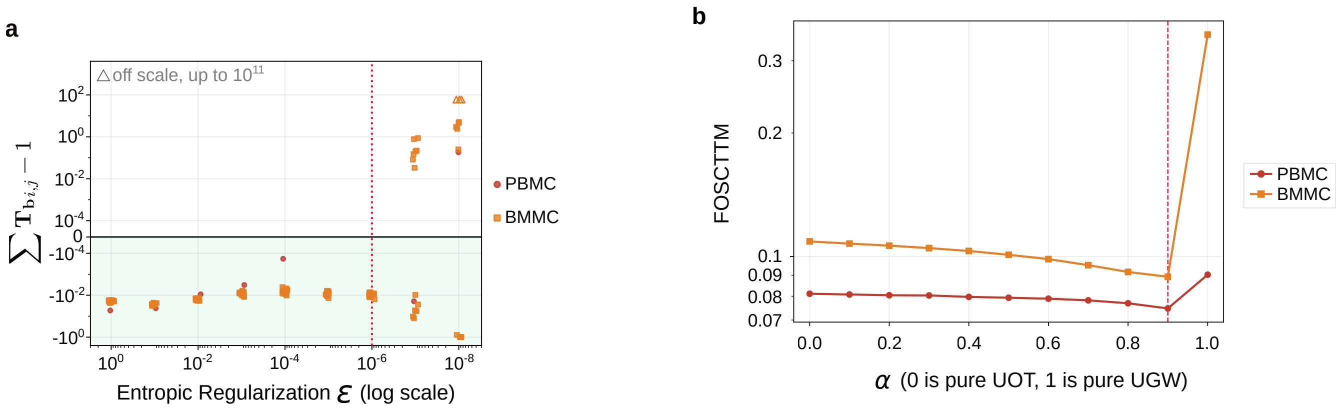


**Figure S1.** **a)** Scatter plot showing the mass deviation ($\sum_{ij} \mathbf{T}_{\mathbf{b}ij} - 1$, y-axis) versus the entropic regularization $\varepsilon$ (x-axis). Positive mass deviation indicates a numerically degenerate solver; $\varepsilon = 10^{-6}$ is the smallest value (faster solver) at which the solutions remain stable. **b)** FOSCTTM (y-axis) versus the $\alpha$ (x-axis) on the PBMC and BMMC validation datasets. Lower FOSCTTM indicates better alignment, and $\alpha = 0.9$ is consistently optimal across both datasets.

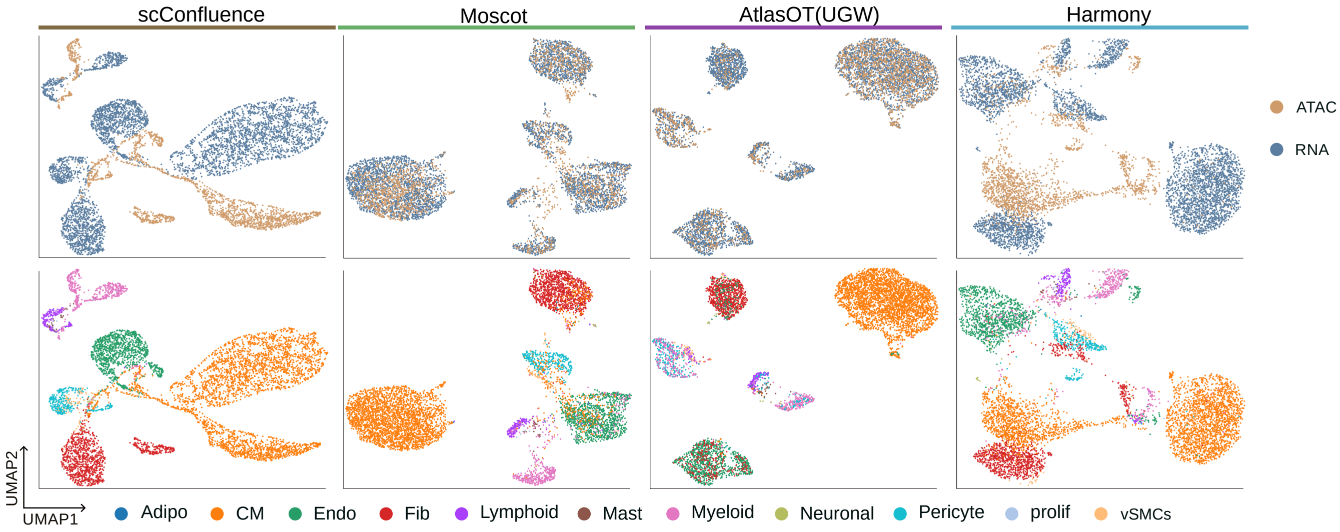


**Figure S2.** Example UMAPs with integrated datasets and cell type annotation for a selected sample RZ_P11 from Kuppe-2022 for algorithms not shown in the main manuscript.

a.

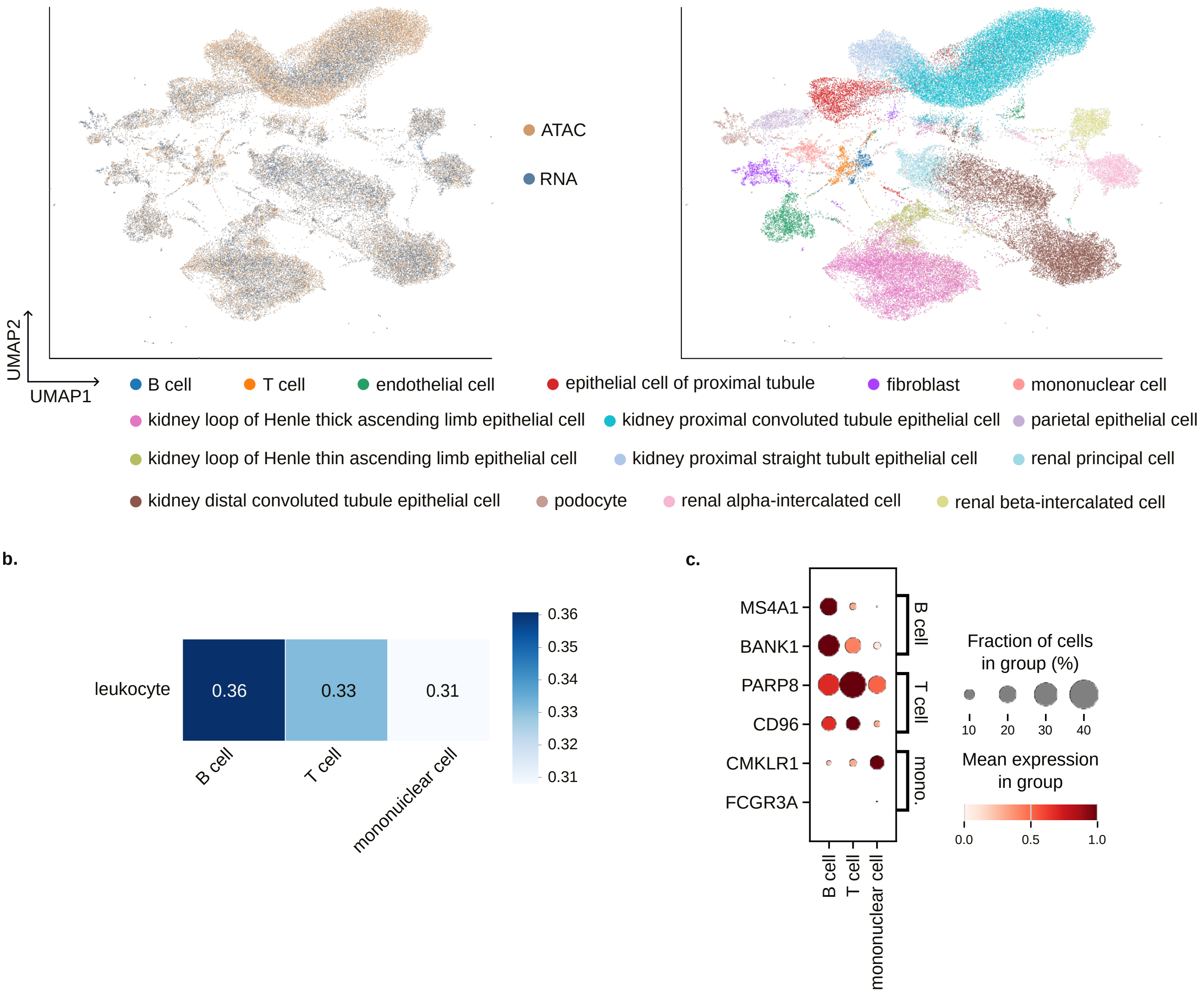


**Figure S3.** **a)** UMAP with all samples from the Wilson-2022 kidney data showing the two modalities (left) and cell types after transfer (right) as estimated by AtlasOT. **b)** The confusion matrix showing how cell labels change from the original (y-axis) vs. transferred (x-axis) cell types. **c)** The marker gene dot plot for RNA leukocyte cells transferred cell types.

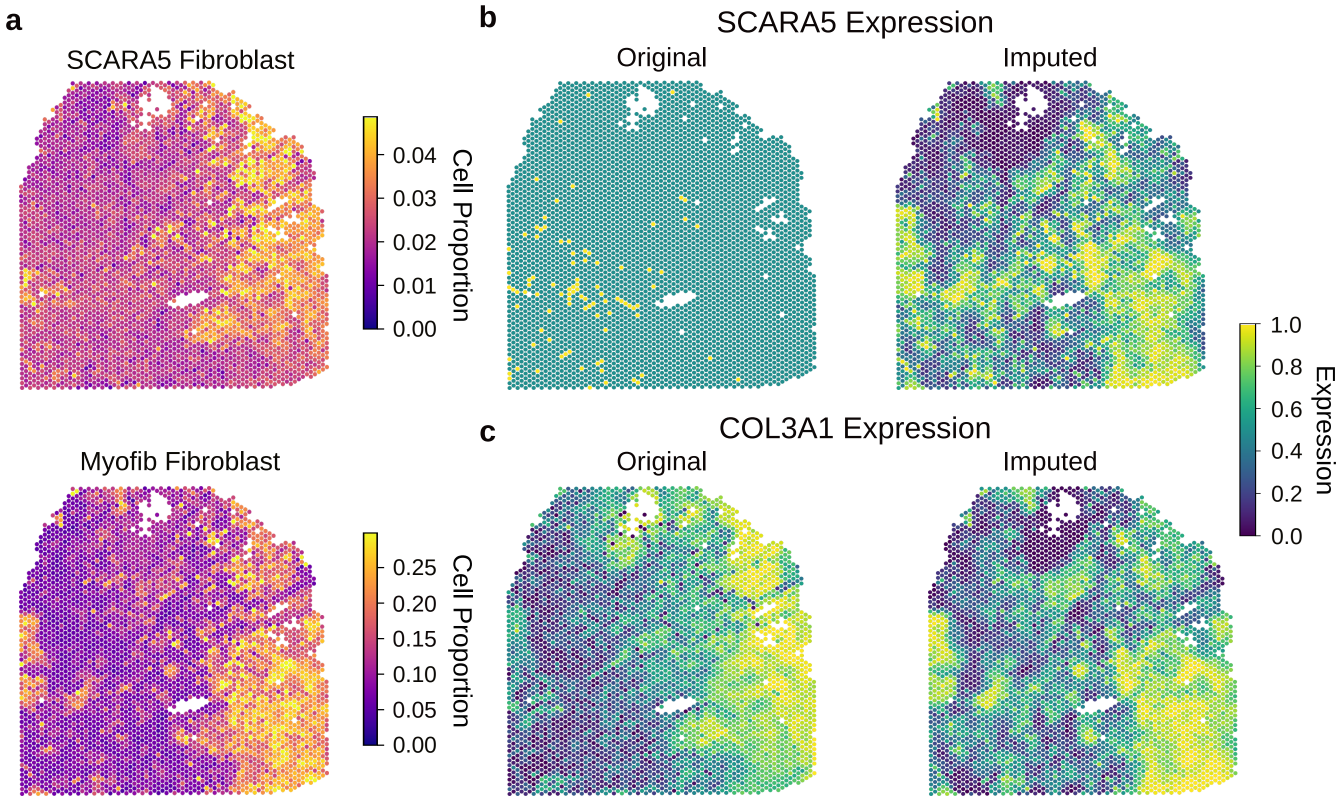


**Figure S4.** **a)** Deconvolution abundance of fibroblast cells subtypes: Myofib and SCARA+ Fib. on spatial spots in an ischaemic (GT_IZ_P9) sample. **b)** Expression values of the original and imputed SCARA5 gene in an ischaemic (GT_IZ_P9) sample. **c)** Expression values of the original and imputed COL3A1 gene in an ischaemic (GT_IZ_P9) sample.

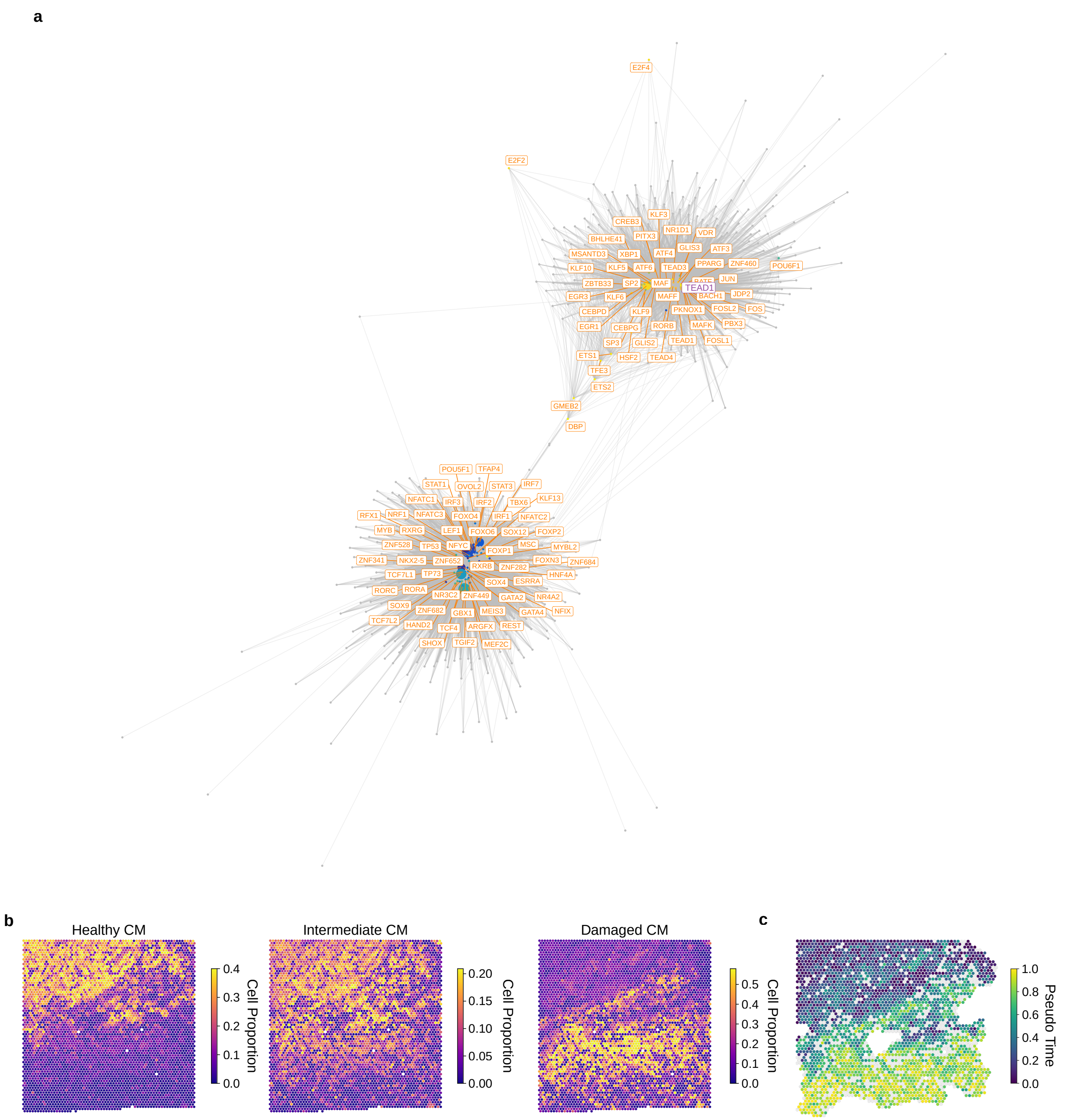


**Figure S5.** **a)** Inferred TF–gene regulatory network (GRN) from the mapped snATAC-snRNA-seq data. TF nodes are coloured by trajectory time point (from blue to yellow) and target-gene nodes are grey; node size scales with network importance; edges denote TF–gene correlations. The representative TF TEAD1 is highlighted in purple. **b)** Deconvolution abundance of cardiomyocytes from healthy to damaged areas in the RZ_BZ_P3 border zone sample. **c)** Spatial pseudotime heatmap of the cardiomyocyte (CM) domain in a human myocardial infarction heart (RZ_BZ_P3), coloured by rank-normalised CM RNA diffusion pseudotime (early to late).

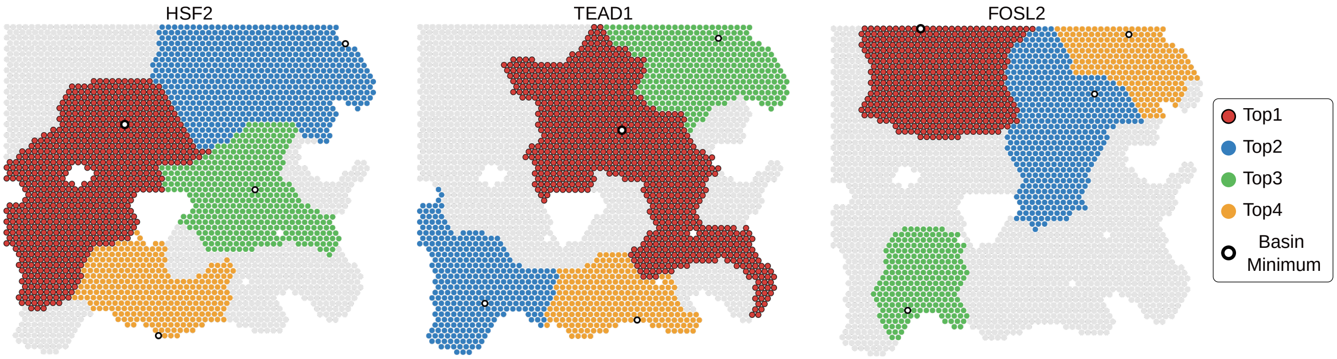


**Figure S6.** The four largest basins of the chromatin flow field for HSF2, TEAD1 and FOSL2 of sample RZ_BZ_P3. Each basin drains to its basin minimum.
